# Bidirectional Crosstalk Between the SAGA Complex Deubiquitinase Module and the Molecular Circadian Clock

**DOI:** 10.64898/2025.12.14.694213

**Authors:** Kara M Costanzo, Zoya Bangash, Cj Talton, Jamille Chamon, Alisa Voyvevidko, Asia Flint, Wei-Ling Tsou, Jin-Yuan Fan, Jeffrey Price, Ryan D Mohan

## Abstract

The Spt-Ada-Gcn5-acetyltransferase (SAGA) complex is a highly conserved chromatin-modifying transcriptional coactivator that regulates gene expression through both transcriptional and post-translational mechanisms. In addition to histone acetyltransferase activity, SAGA harbors a deubiquitinase module (DUBm) that influences protein stability, localization, and function by removal of ubiquitin. Although SAGA’s roles in transcription are well characterized, how its enzymatic modules are dynamically regulated to coordinate its divergent activities remains less known. Here, we identify a bidirectional relationship between the molecular circadian clock and the SAGA DUBm. Our results indicate that circadian rhythms influence DUBm activities and expression, while the DUBm in turn shapes the timing and stability of core clock proteins and transcripts, positioning the DUBm as an important mediator of the molecular circadian clock.

## Introduction

The discovery of a direct connection between transcriptional activation and changes in chromatin structure via post-translational modifications such as histone acetylation brought about a major advancement in our understanding of gene regulation in eukaryotes. The Spt-Ada-Gcn5-acetyltransferase (SAGA) chromatin-modifying complex is a highly conserved, approximately 1.8 MDa transcriptional coactivator complex comprised of ∼20 subunits arranged within a modular fashion (Cloud et al., 2019; Cornelio-Parra, Goswami, Costanzo, Morales-Sosa, & Mohan, 2021; Grant, Winston, & Berger, 2021; Lee, Swanson, Florens, Washburn, & Workman, 2009). SAGA utilizes two enzymatic activities to coordinate gene expression: its enzymatic histone-acetyltransferase activity, housed within the GCN5/Pcaf subunit, and its deubiquitinase activity, associated with the yUbp8/dnon-stop/hUsp22 subunit (Grant et al., 2021; Lee, Florens, Swanson, Washburn, & Workman, 2005; Sterner et al., 1999). Mutations in SAGA’s deubiquitinase module (DUBm) results to the neurodegenerative disease Spinocerebellar ataxia type 7 (SCA7), caused by CAG triplet expansion of the *Ataxin-7* (*Atxn7*) gene, leading to polyglutamine expansion in the expressed Ataxin-7 protein (Cornelio-Parra et al., 2021; David et al., 1997; Goswami et al., 2022).

The size of CAG repeats ranges from 4 to 18 in normal alleles and from 36 to 460 repeats in expanded alleles (David et al., 1997; Goswami et al., 2022; La Spada, 1993). In SCA7, the length of the polyglutamine tract affects both age of disease onset and severity of disease. A longer polyglutamine tract is correlated with an earlier age of onset and more severe symptoms (Garden & La Spada, 2008; Goswami et al., 2022; La Spada, 1993). In *Drosophila*, loss of *Atxn7* leads to a phenotype similar to overexpression of the poly-Q-expanded amino terminal truncation of human Atxn7 (Latouche et al., 2007) – reduced life span, reduced mobility, and retinal degeneration (Mohan et al., 2014).

Although SAGA’s roles in transcription have been characterized, less is known about how the individual modules and enzymatic activities of SAGA are coordinated and cause such specific disease phenotypes. For organisms to adapt and survive, biological and biochemical events must be precisely coordinated to occur at specific times of day (Roenneberg & Merrow, 2016). This temporal regulation is governed by internal circadian clocks and transcriptional-translational feedback loops aligned with Earth’s 24-hour light/dark cycle that orchestrate daily physiological and molecular processes (Fuhr, Abreu, Pett, & Relogio, 2015; Roenneberg & Merrow, 2016; Takahashi, 2017). These clocks are primarily located in the brain, and crosstalk between the circadian molecular clock and other key intracellular pathways and regulators further shape mammalian physiology (Fagiani et al., 2022).

Chromatin-modifying and transcriptional coactivator complexes play critical roles in maintaining these regulatory loops. Amongst them, the SAGA complex was one of the first identified transcriptional coactivators consisting of modular components with diverse transcriptional regulatory functions (Grant et al., 2021). The deubiquitinase module of SAGA has established activities towards histone H2B (H2B), histone H2A (H2A), shelterin, and FUSE-binding protein 1 (FBP1) (Atanassov & Dent, 2011; Daniel et al., 2004; Henry et al., 2003; Weake et al., 2008).

Across species, DUBm can dissociate from SAGA and remain enzymatically active (Lim, Kwak, Kim, & Lee, 2013), supporting its independent regulatory roles (X. Li et al., 2017). Loss of *Atxn7* releases the DUBm from SAGA and is lethal, though lethality can be rescued by reducing *non-stop (not)* gene copy number, suggesting a class of SAGA unbound non-stop target(s) that may contribute to cytotoxicity (Mohan et al., 2014).

Traditionally, transcriptional coactivator complexes have been viewed as static assemblies recruited to chromatin as fixed monolithic complexes. However, evidence from our lab demonstrates that the DUBm association with SAGA is dynamic. Using *Drosophila* as a model, our findings suggest that SAGA’s DUB module and its enzymatic activities have a bidirectional relationship with the circadian clock. Circadian rhythms influence post-translational control of DUBm protein levels, while the DUBm in turn affects the stability of core clock proteins and transcripts. Together, these findings support a model in which a bidirectional relationship between the circadian clock and SAGA DUBm components together maintain post-translational modification dynamics and circadian rhythmicity.

## Results

### SAGA DUBm genes are circadian across tissues

To determine whether components of the SAGA complex are transcriptionally regulated by the circadian clock, we analyzed publicly available RNA-sequencing (RNA-seq) datasets in mice across multiple tissues and circadian time points. We compared expression profiles of genes encoding the SAGA deubiquitinase module, including *not* and *Atxn7*, with those genes encoding core structural and enzymatic components of the larger SAGA complex, such as *Ada3* and *Taf6*. As shown in figure 1, three of the four DUBm subunit transcripts exhibit circadian oscillations over the 24-hour cycle, indicating that these genes are expressed under circadian control (Figure 1B). In contrast, many transcripts encoding the core SAGA machinery and other modules such as *Ada3, Taf5*, and *Supt3* do not display rhythmic cycling as seen with DUBm. This was observed across multiple tissues in mice.

**Figure 1.**
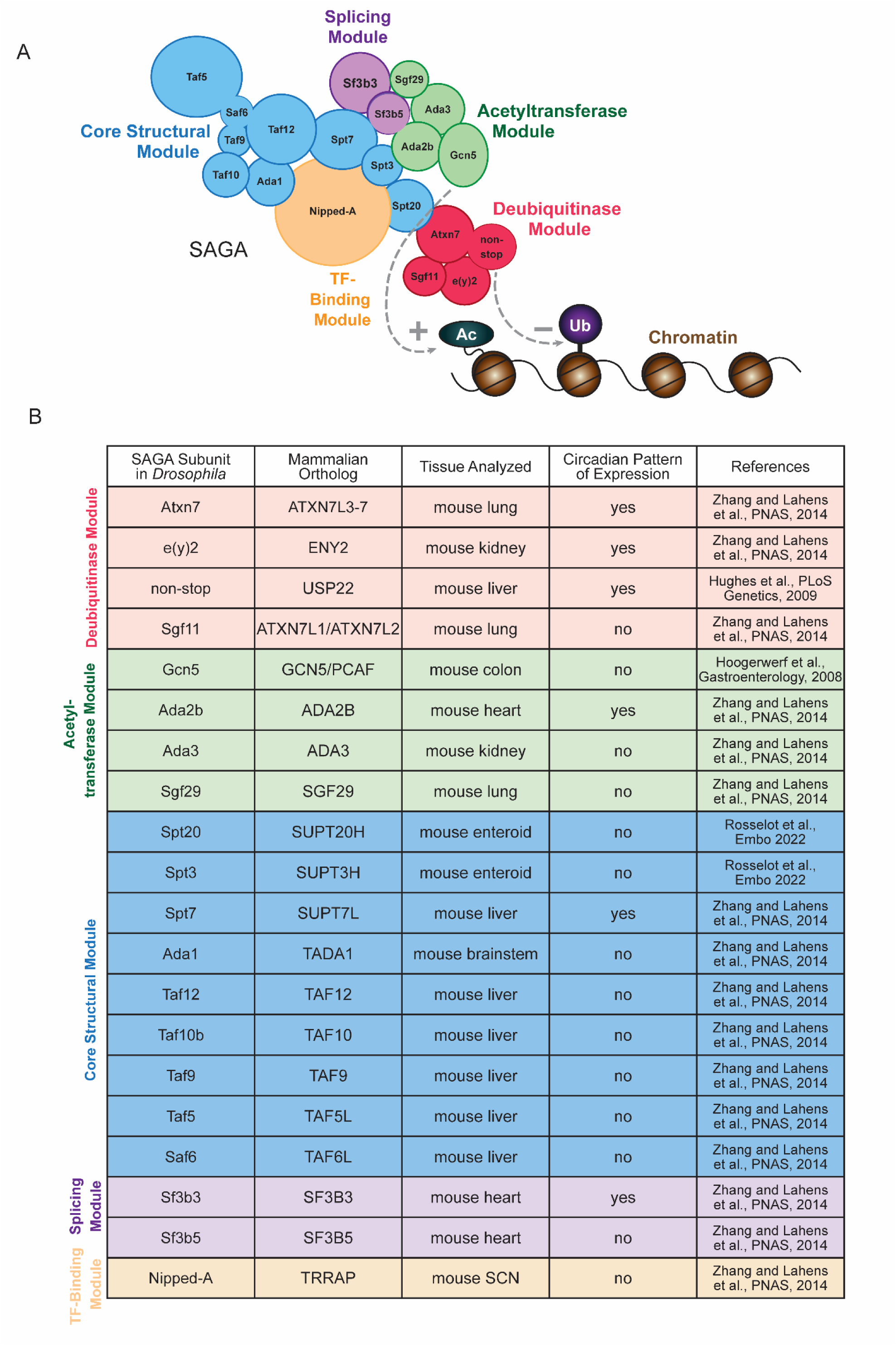
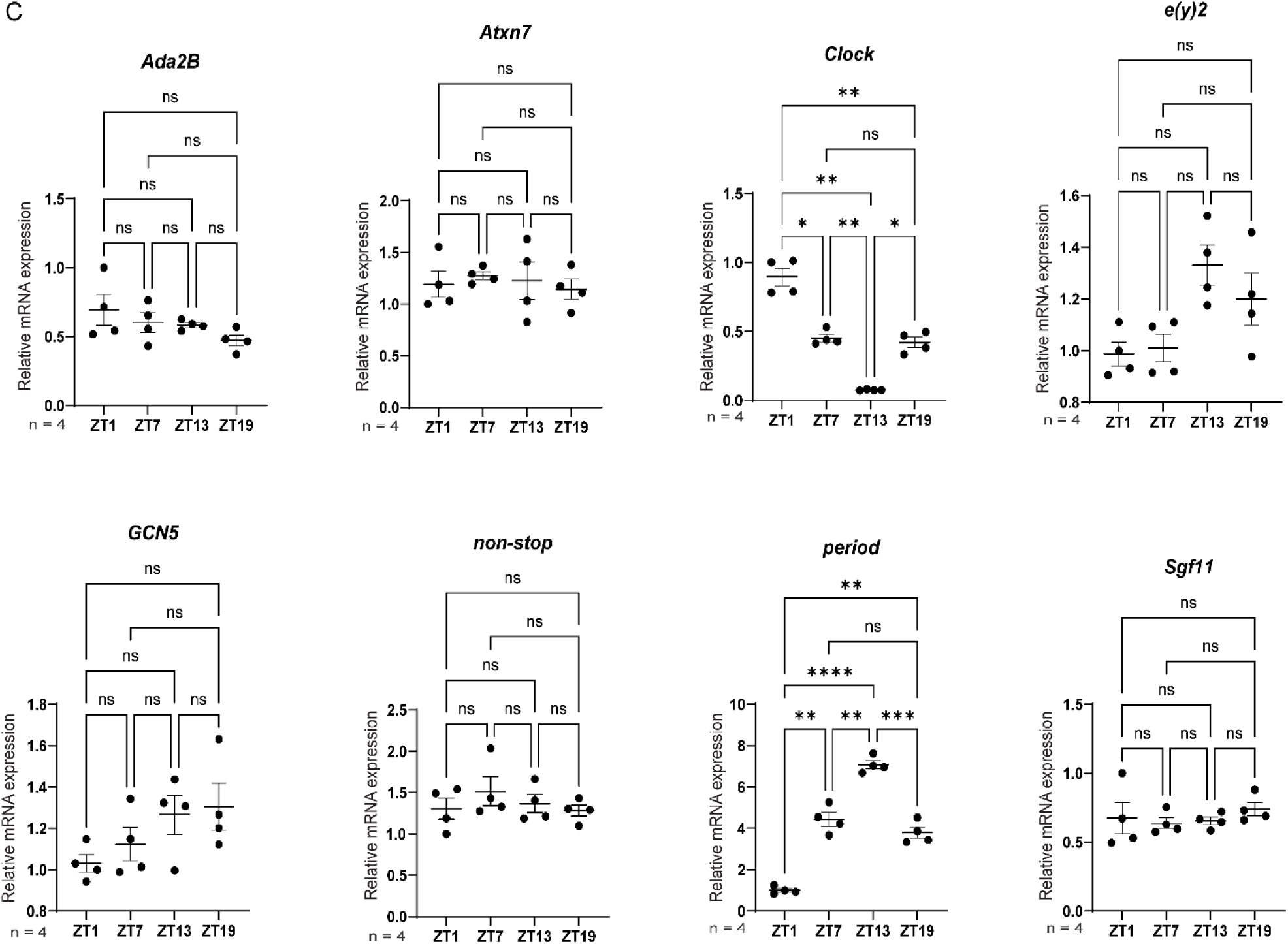
Modular circadian regulation of the SAGA deubiquitinase module. (A) Schematic representation of the SAGA complex, illustrating its major functional modules, including the deubiquitinase module (DUBm), histone acetyltransferase (HAT) module, core structural components, and transcription factor–interacting subunits. (B) Mouse RNA-seq analysis across multiple tissues reveals rhythmic, circadian oscillations in three of four DUBm transcripts, including *not* and *Atxn7*, while transcripts encoding core SAGA components (*Ada3*, *Taf5*, *Supt3*) are non-rhythmic. (C) qRT-PCR shows no significant circadian oscillations in DUBm transcript levels in *Drosophila* whole-head extracts, suggesting species-specific differences or post-translational regulation in flies. *p < 0.05, **p < 0.005, ***p < 0.0005, ****p < 0.0001, ns is not significant. *Period* (p < 0.0001) and *Clock* (p < 0.0001) serve as a circadian control and show robust and significant circadian oscillations. Welch ANOVA with Brown-Forsythe post-hoc was performed, and error bars represent standard error of the mean (SEM).

This distinction suggests that circadian regulation within SAGA is modular, selectively influencing the DUBm rather than the complex as a whole. Such targeted rhythmic regulation is likely to be physiologically significant. By coupling DUBm expression to circadian time, cells may achieve a dynamic balance between the DUBm’s functions inside and outside of the SAGA complex. Unbiased proteomic screens have identified diverse DUBm interaction partners, highlighting a broad and expanding functional network. Temporal control of DUBm abundance may therefore serve as a mechanism to coordinate its many activities with daily cellular and organismal rhythms.

We next examined whether this circadian regulation is conserved in flies at the transcript level. In circadian studies, Zeitgeber time (ZT) is often used as a standardized 24-hour notation of the phase in an entrained circadian cycle in which ZT0 indicates the beginning of the day, or the light phase, and ZT12 is the beginning of the night, or dark phase (Vitaterna, Takahashi, & Turek, 2001). To directly evaluate whether SAGA components are subject to circadian transcriptional regulation, we performed qRT-PCR on heads of circadian-entrained *Drosophila*. Flies were entrained in a 12:12 light-dark cycle for three to five days and collected at the following four time points: ZT1, representing 1 hour after lights go on; ZT7, representing 7 hours after lights go on; ZT13, representing 1 hour after lights go off; and ZT19, representing 7 hours after lights go off.

In contrast to the mouse datasets, qRT-PCR analysis of SAGA DUBm genes in *Drosophila* whole-head extracts did not show significant circadian oscillations at the transcript level (Figure1C). *Clock* and *period* served as a control and displayed typical and robust circadian mRNA expressions. One possibility for this result is that circadian regulation occurs post-translationally in flies, affecting DUBm protein stability, localization, or activity rather than transcript abundance. Another consideration is that whole-head measurements may mask tissue-specific rhythms, For example, opposing patterns in eye versus brain tissues. These findings prompted us to next examine whether circadian control occurs at the protein level, rather than the transcript level.

### SAGA DUBm proteins non-stop and Atxn7 fluctuate throughout the circadian cycle in a per-dependent manner

Proper coordination and expression of the SAGA DUBm is critical for biological homeostasis, as its mutation leads to aberrant gene expression, defective non-homologous end joining, decreased immune response, reduced replicative lifespan, neurodegeneration, blindness, tumorigenesis, and cancer (Bonnet et al., 2014; Furrer et al., 2011; Glinsky, Berezovska, & Glinskii, 2005; Lang et al., 2011; C. Li et al., 2018; McCormick et al., 2014; Mohan et al., 2014; Weake et al., 2008). In addition, reports from patients with SCA7 include disturbances in circadian rhythms, such as sleep disorders (insomnia, snoring, and restless legs syndrome) and hyperkinetic movement disorders (chorea and myokymia), which are more frequent in patients with early onset SCA7 than in patients with adult onset (Velazquez-Perez et al., 2015).

Based on findings from our lab and others, we propose circadian regulation serves as a mechanism that coordinates DUBm dynamics, and Atxn7 poly-Q expansion conditions may hinder non-stop’s circadian rhythmicity, altering gene expression and resulting in SCA7 disease phenotypes. As a circadian control, wild-type flies are compared to flies barring a loss of circadian rhythm via a *per* loss of function allele (*per^0^*).

We immunoblotted protein extracts prepared from wild-type and *per^0^* fly heads taken at each of the four ZTs and found that non-stop protein levels significantly fluctuate in wild type-flies throughout the day. Between the ZT with the lowest expression and the ZT with the highest expression, there was a nearly three and a half-fold increase in not (Figure 2A). *Per^0^*flies did not show fluctuations in expression, with relatively consistent non-stop protein levels throughout all timepoints (Figure 2B).

**Figure 2.**
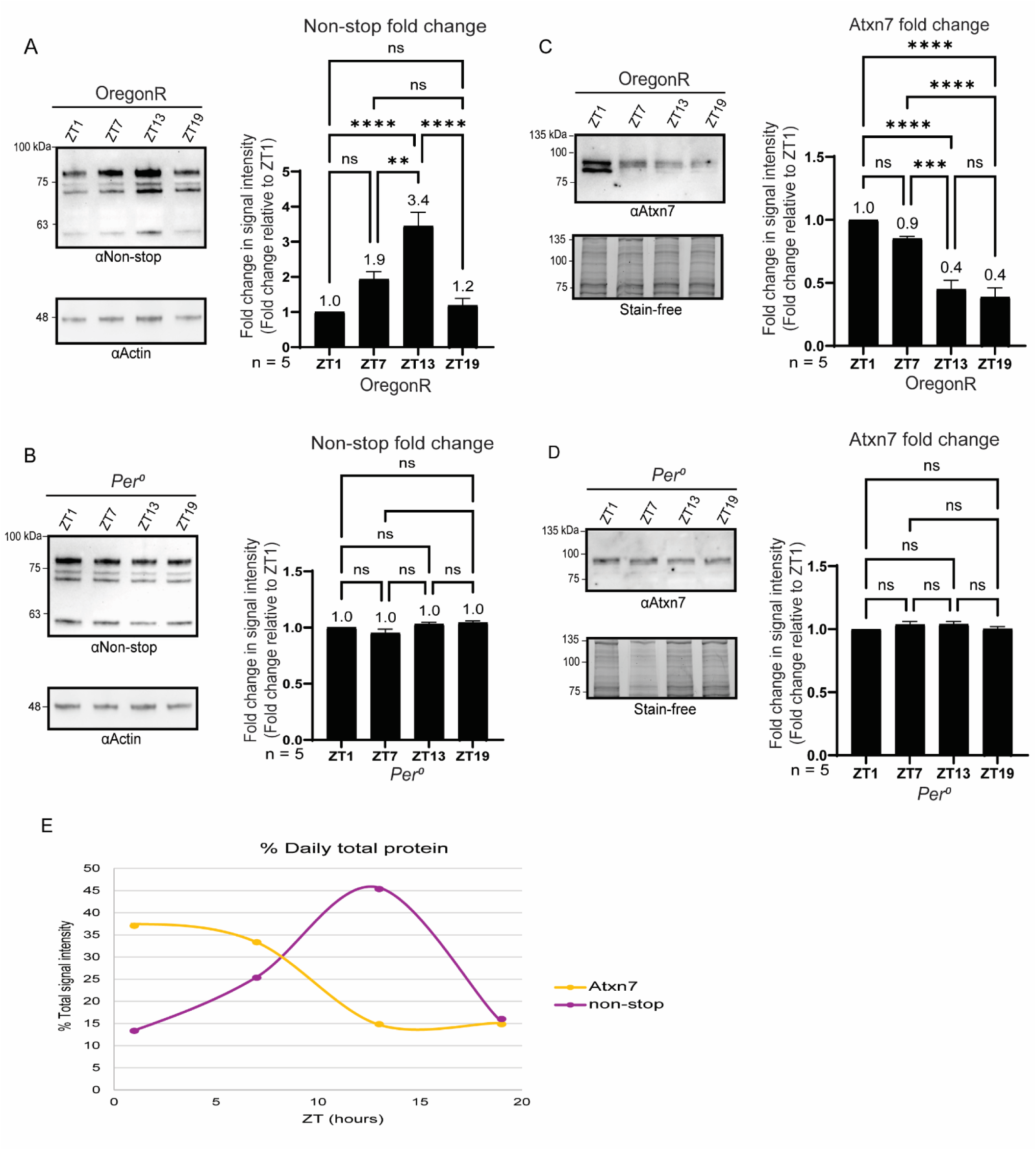
SAGA DUBm proteins non-stop and Atxn7 are expressed in a circadian pattern. Immunoblotting against anti-non-stop in circadian entrained flies reveals a circadian pattern of protein expression in wild-type flies (A) that was abolished in *per^0^* flies (B). The levels of non-stop quantified from at least three biological repeats are shown in the graphs to the right. Statistical analysis was performed using One-way ANOVA followed by Tukey’s multiple comparison test, with error bars representing standard error of the mean (SEM); significant differences in non-stop levels were detected between ZT1 and ZT7 (p=0.01), ZT1 and ZT13 (p=0.001), and ZT13 and ZT19 (p=0.005). (*C*) Immunoblotting against anti-*Atxn7* in circadian entrained flies reveals circadian regulation in wild-type background that was abolished in *per^0^* flies (D). The levels of Atxn7 quantified from at least three biological repeats are shown in the graphs to the right. Statistical analysis was performed using One-way ANOVA followed by Tukey’s multiple comparison test, with error bars representing standard error of the mean (SEM); significant differences in Atxn7 levels were detected between all pairs of ZTs are indicated. *p < 0.05, **p < 0.005, ***p < 0.0005, ****p < 0.0001, ns is not significant. (*E*) Averages across ZTs were calculated as a percentage of daily total and plotted across circadian time.

Circadian protein expression was also observed for Atxn7 in wild-type flies (Figure 2C), whereas *per^0^* flies did not show significant fluctuations in expression and displayed relatively consistent Atxn7 protein levels throughout all timepoints (Figure 2D). Between the ZT with the lowest expression and the ZT with the highest expression, there was a two and a half-fold increase in Atxn7 protein. Averages across ZTs were calculated as a percentage of daily total and plotted across circadian time, demonstrating times where Atxn7 and non-stop expression levels are essentially inverse (ZT1, ZT13) (Figure 2E).

### Circadian SAGA model of control

Loss of *Atxn7* causes release of not from the SAGA complex and results in lethality. However, reducing the gene copy number of *non-stop* in flies with loss of *Atxn7* rescues lethality (Cloud et al., 2019). This suggests the existence of non-stop target(s) that bind more strongly to non-stop when Atxn7 is absent or mutated. Additionally, it raises the possibility that some of these targets may contribute to the toxicity associated with the loss or polyglutamine expansion of Atxn7. We propose a model in which the SAGA complex is dynamically regulated by the circadian clock, allowing its deubiquitinase module to cycle between SAGA-bound and SAGA-independent functions over the course of the day (Figure 3).

**Figure 3.**
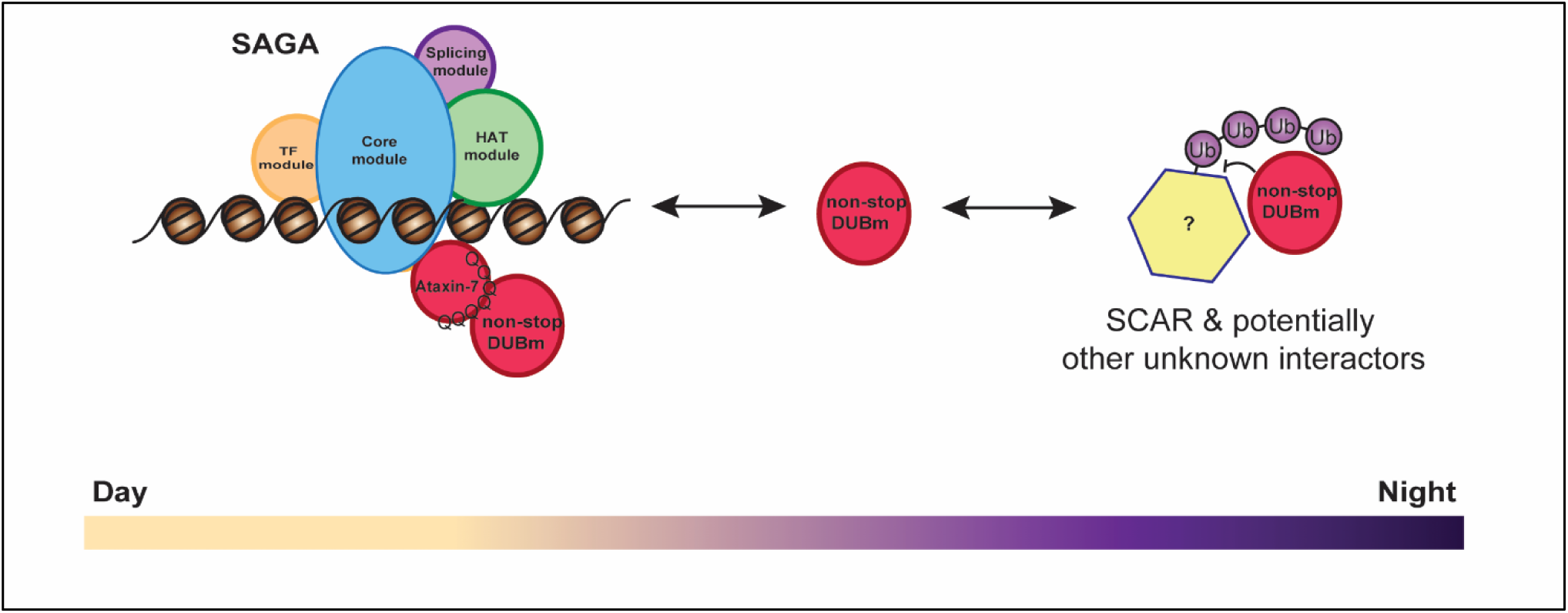
Proposed model for circadian regulation of the SAGA complex and its DUBm. Conceptual model illustrating how circadian timing may influence DUBm associations with SAGA and its independent substrates. This rhythmic association provides a potential mechanism for coordinating DUBm activity with circadian and cellular physiological demands.

### Non-stop deubiquitinase substrate SCAR is expressed in a circadian pattern

The mechanisms by which the SAGA DUBm interacts with complexes outside of the core SAGA complex are not fully understood. If non-stop changes expression and interaction partners, this would likely be reflected by changes in non-stop substrate levels. Consistent findings from different laboratories have shown that DUBm can function as an independent module (Cloud et al., 2019; Helmlinger et al., 2004; Lang et al., 2011; Lim et al., 2013). In *Drosophila*, the DUBm can separate from SAGA and interact with the WAVE regulatory complex (WRC) through conserved WRC interacting receptor sequences (WIRS) which decorate the amino and carboxy ends of non-stop/USP22 (Cloud et al., 2019; Cornelio-Parra et al., 2021). There, non-stop counters ubiquitination and proteolytic degradation of WAVE, also known as SCAR in *Drosophila*, the active subunit of the WRC (Cornelio-Parra et al., 2021).

SCAR functions within WRC to activate the Actin-related protein 2/3 (Arp2/3) complex and facilitate branching of F-actin filaments (Kurisu & Takenawa, 2009). In *Drosophila*, SCAR is particularly important for cytoplasmic organization in the blastoderm, for egg chamber structure during oogenesis, axon development in the central nervous system, and adult eye morphology (Hudson & Cooley, 2002; Zallen et al., 2002).

SCAR is regulated through a balance of ubiquitination and deubiquitination (Kunda, Craig, Dominguez, & Baum, 2003), and we found up to 70% of SCAR levels to be regulated by non-stop (Cloud et al., 2019), emphasizing the importance of non-stop in maintaining its stability. SAGA, WRC, and Arp2/3 complexes are all critical for proper central nervous system function, making the regulatory intersection for these complexes intriguing for further study (Chou & Wang, 2016; Dumpich, Mannherz, & Theiss, 2015; Kunda et al., 2003; G. Meyer & Feldman, 2002; Mohan et al., 2014; Zallen et al., 2002).

To demonstrate that non-stop is a primary regulator of SCAR levels in *Drosophila*, we immunoblotted extracts prepared from wild-type and non-stop loss-of-function mutant (*non-stop^0206^)* whole larvae. Our immunoblots consistently show that loss of *non-stop* causes a five-fold decrease in SCAR protein levels (Figure 4A). Having established non-stop is a primary regulator of SCAR, we sought to explore whether SCAR is circadian and examined SCAR expression in heads of circadian-entrained *Drosophila* as explained above. Immunoblot analysis revealed that SCAR oscillates in a circadian pattern similar to non-stop, with a more than two and a half-fold increase in SCAR from the lowest to highest ZT (Figure 4B), an effect that was not observed in *per^0^* mutants (Figure 4C). Averages across ZTs were calculated as a percentage of daily total and plotted across circadian time alongside Atxn7 and non-stop (Figure 4D).

**Figure 4.**
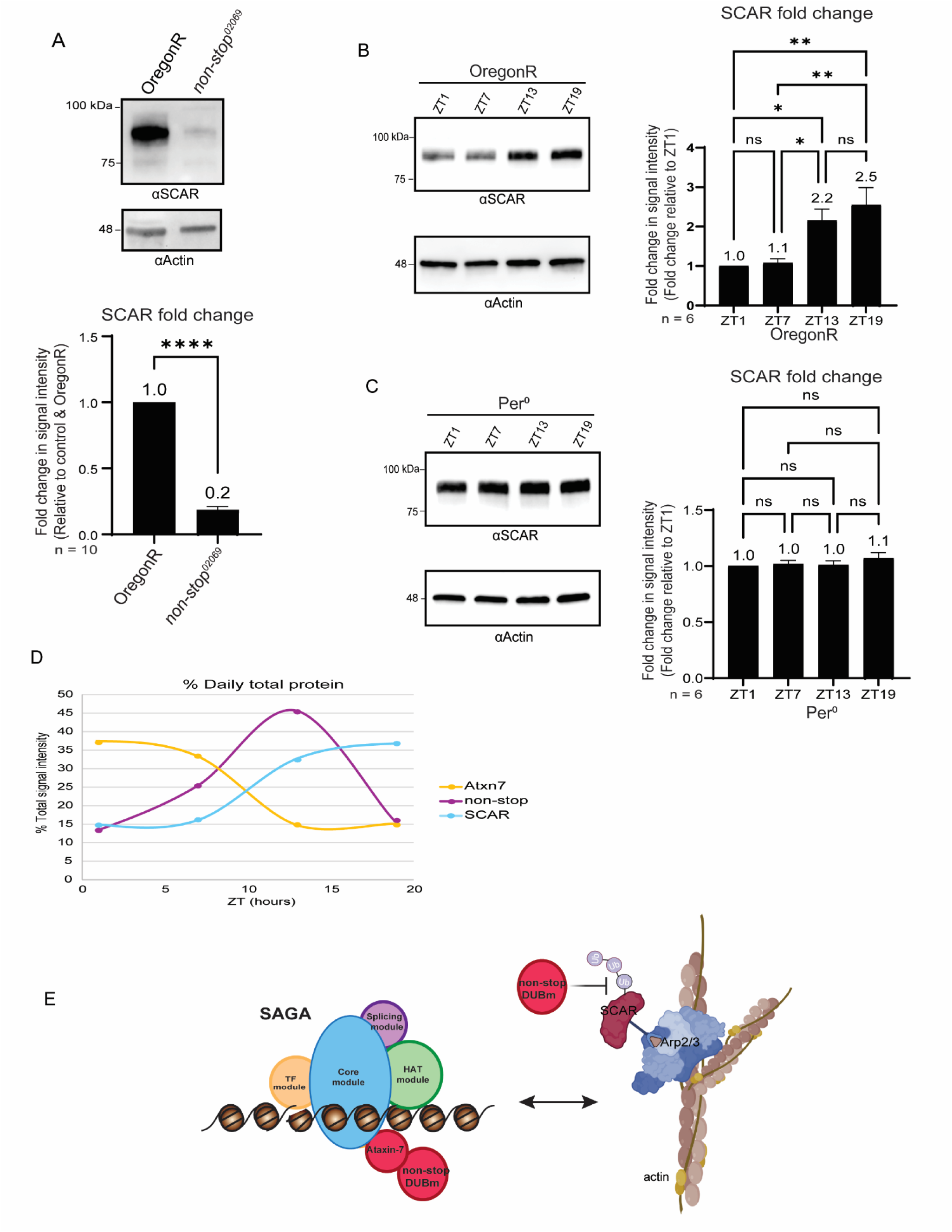
WRC subunit SCAR is expressed in a circadian pattern. (A) Non-stop is a primary regulator of SCAR, and loss of *non-stop* causes a five-fold decrease in SCAR protein levels. The levels of SCAR quantified from at least three biological repeats are shown in the graph below. Statistical analysis was performed using a paired t-test with error bars representing standard error of the mean (SEM) (*p < 0.05, **p < 0.005, ***p < 0.0005, ****p < 0.0001). (B) Immunoblotting against anti-SCAR in circadian entrained flies reveals circadian expression patterns in the wild-type background that are abolished in *per^0^* flies (C). The levels of SCAR quantified from at least three biological repeats are shown in the graphs to the right. Statistical analysis was performed using one-way ANOVA followed by Tukey’s multiple comparison test, with error bars representing standard error of the mean (SEM). *p < 0.05, **p < 0.005, ***p < 0.0005, ****p < 0.0001, ns is not significant. (D) Averages across ZTs were calculated as a percentage of daily total and plotted across circadian times. (E) Hypothetical model for dynamic DUBm SAGA-WRC associations mediated through non-stop.

Taken together, we created a model of balanced circadian non-stop enzymatic activity with SAGA and WRC (Fig 4E). The concept is that the circadian clock imposes rhythmic control on the DUBm, but not on the full SAGA complex. Such regulations may serve to balance DUBm functions within SAGA (i.e. H2B deubiquitination and transcriptional regulation) and outside of SAGA (i.e. SCAR deubiquitination and stabilization), ensuring that these activities are temporally aligned with daily physiological rhythms.

### Non-stop and Atxn7 exhibit distinct and dynamic localization patterns in the brain over circadian time

To investigate whether components of the SAGA DUBm undergo circadian-dependent spatial reorganization, we examined the subcellular localization of not and Atxn7 across the circadian cycle. Brains from entrained *Drosophila* were immunoassayed at four ZTs to assess whether these proteins exhibit time-of-day-dependent changes in expression or compartmentalization. Maximum intensity projections generated from Z-stack confocal images were used for visualization, providing a qualitative overview of localization patterns while acknowledging that this method emphasizes the brightest signals in each channel rather than uniformly sampling volumetric data (Figure 5).

**Figure 5.**
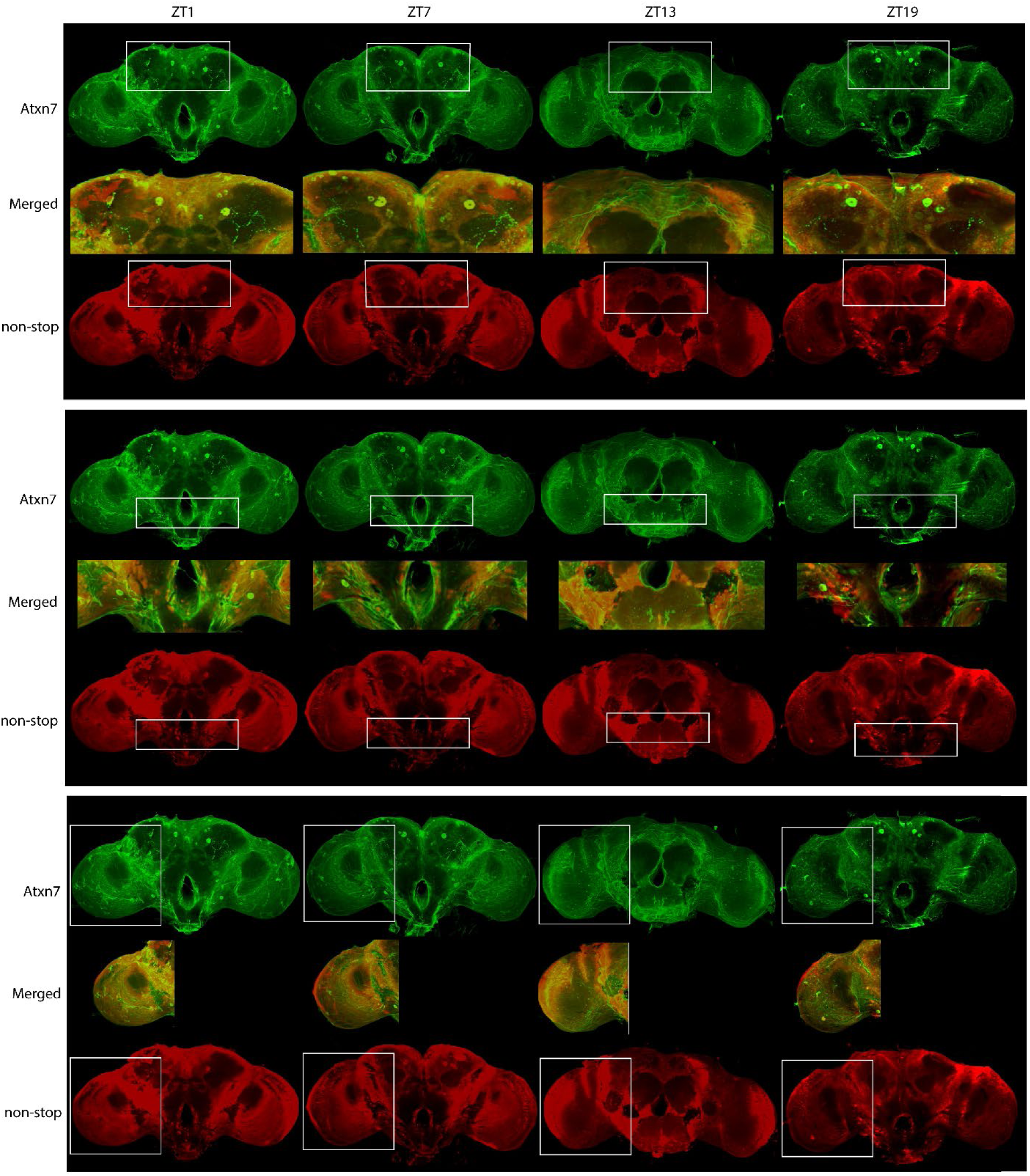
Non-stop and Atxn7 exhibit distinct and dynamic localization patterns over circadian time. Brains were dissected and stained at four Zeitgeber times (ZT1, ZT7, ZT13, and ZT19) to assess time-of-day spatial distribution of Atxn7 (green) and non-stop (red). Maximum intensity projections were generated from confocal Z-stacks to visualize overall localization patterns.

Arrows indicate cells and notable regions displaying differential expressions or localization across ZTs. Non-stop and Atxn7 display dynamic changes in expression and localization throughout the circadian cycle.

Importantly, our data reveals dynamic changes in localization of both non-stop and Atxn7 across the day in *Drosophila* brains. We observe brain regions and subcellular compartments in which non-stop and Atxn7 do not co-localize (Figure 6). This spatial separation is consistent with growing evidence that non-stop can dissociate from SAGA (Lim et al., 2013; Young et al., 2007) and may shuttle between nuclear and cytoplasmic compartments. It was found that components of the DUBm can associate with other complexes and participate in their roles independently of SAGA (Cloud et al., 2019; Gurskiy et al., 2012). It is possible that non-stop is associating with a separate complex and participating in other regulatory functions at the times and regions, where we do not see Atxn7 and non-stop expressed together.

**Figure 6.**
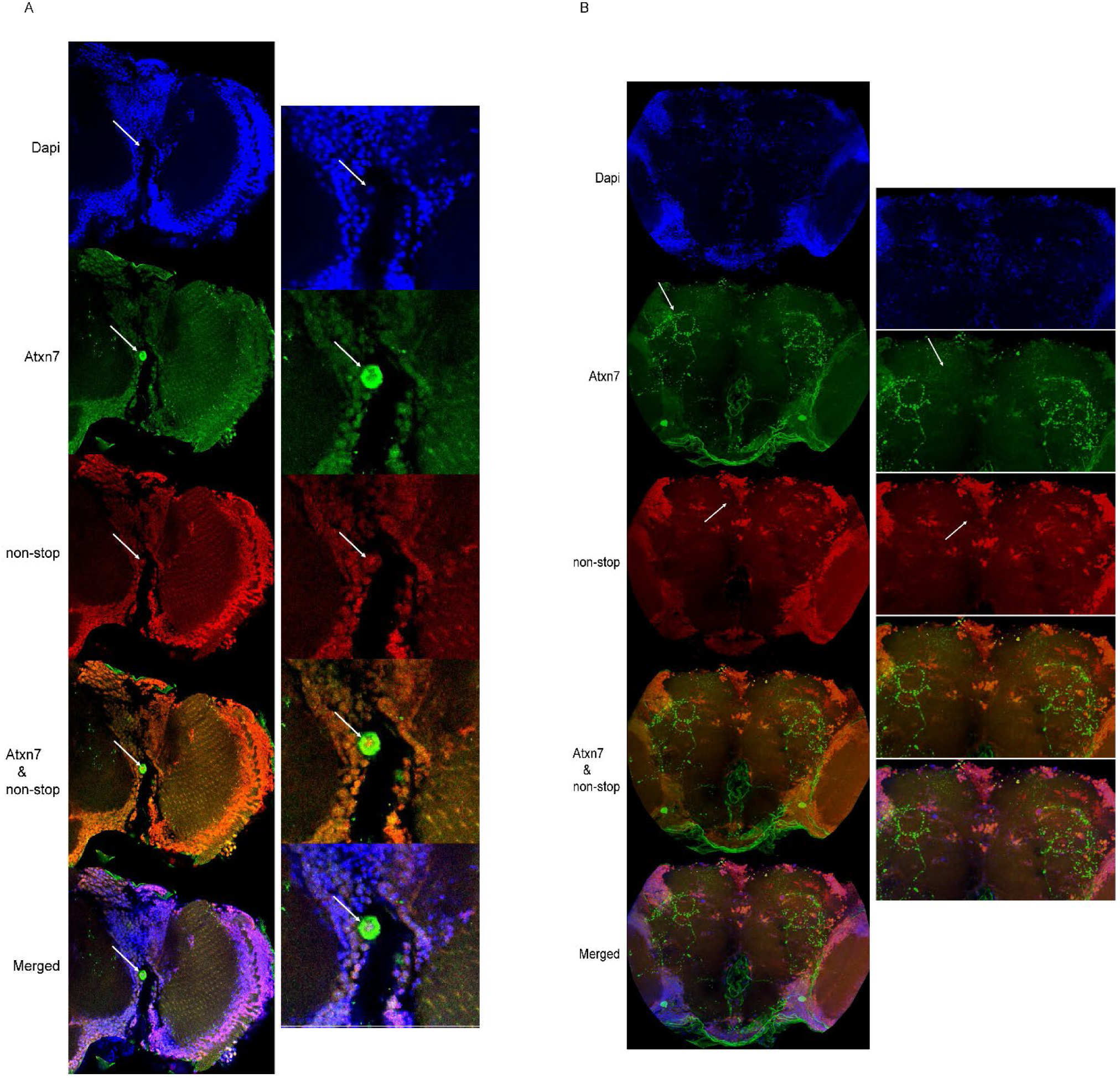
Non-stop and Atxn7 exhibit distinct subcellular localizations in specific cell populations. (A) A cell between the central brain region and optic lobe shows strong nuclear localization of non-stop and predominantly cytoplasmic Atxn7. (B) Additional cells in the central cortex region exhibit the same phenomenon, with non-stop and Atxn7 seen being strongly expressed separately in certain cells and neuronal projections.

### Polyglutamine expansion and loss-of-function mutations in Atxn7 cause disruptions in circadian rhythmicity and period

To determine if Atxn7 affects the circadian clock at the gene or protein level, we did a series of genetic experiments to determine if overexpression of Atxn7, polyglutamine expanded Atxn7, or loss of Atxn7 in clock cells disrupts circadian locomotor profiles and rhythms in flies. We set genetic crosses using RNAi and the UAS-GAL4 system for targeted gene expression to drive different mutations of *Atxn7* into specific cells. *Timeless*-GAL4 is widely used to drive gene expression in the clock cells of the brain, including neurons and glia (Kaneko & Hall, 2000). In contrast, *Pdf*-GAL4 is used to drive expression specifically into pigment dispersing factor (Pdf)-expressing clock cells, which are restricted to the lateral ventral clock neurons (s-LVv and l-LNv) (Renn, Park, Rosbash, Hall, & Taghert, 1999; Sekiguchi, Inoue, Yang, Luo, & Yoshii, 2020).

We found that overexpression of wild-type *Atxn7* using UAS-dcr2>*tim*GAL4 (*tim*GAL4) had minimal effects on locomotor activity profiles (Figure 7C), while overexpressing *poly-Q Atxn7 (UAS-SCA7-100Q)* (Figure 7D), as well as knockdown of *Atxn7* using UAS-*Atxn7* RNAi (Figure 7B) caused significant disruptions in locomotor activity profiles. Data were recorded and analyzed using the method described previously by (Muskus, Preuss, Fan, Bjes, & Price, 2007).

**Figure 7.**
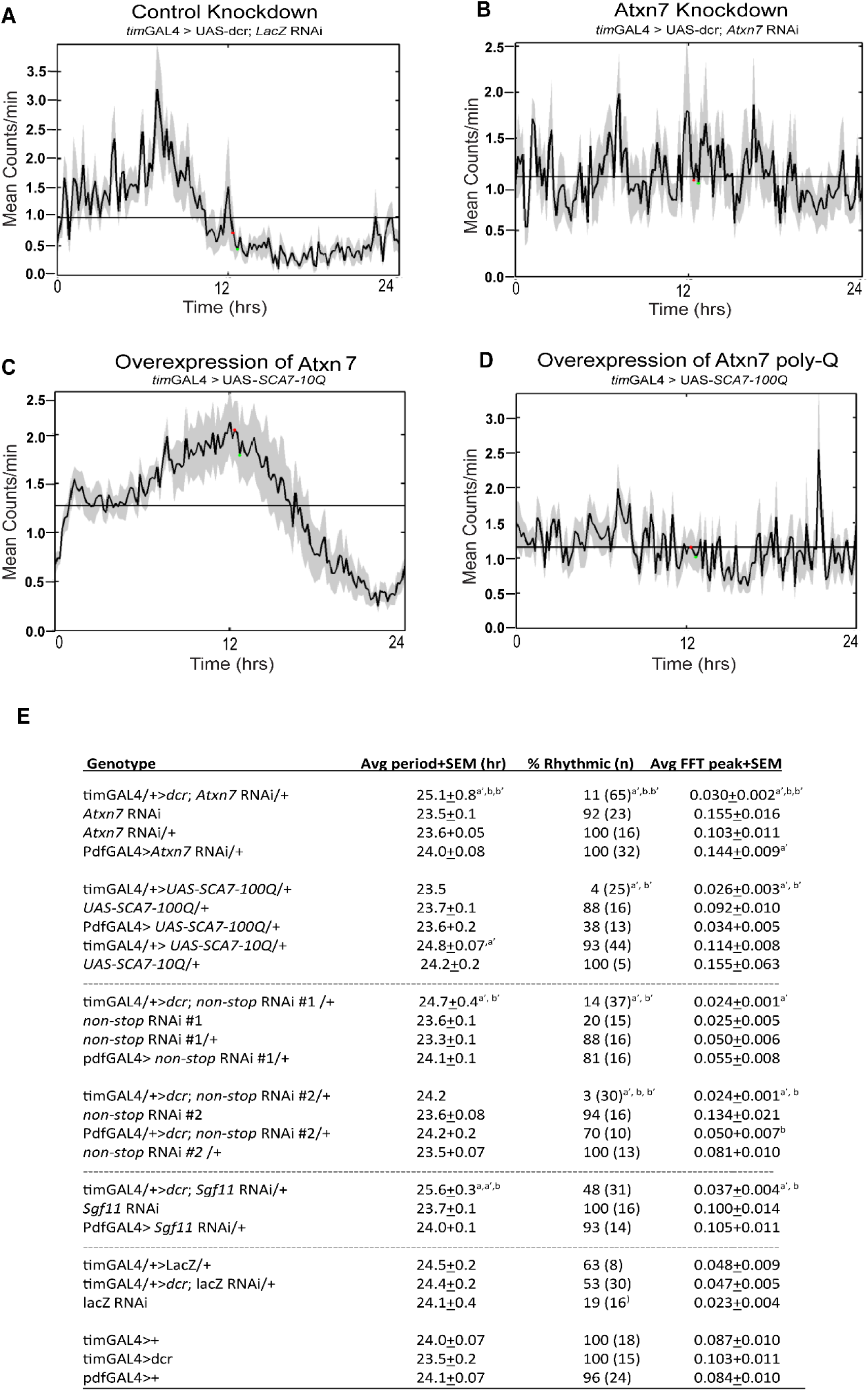
Mutations in Atxn7 cause disruptions in circadian rhythmicity and period. (A) Flies were entrained for at least three days to light/dark and then released into dark/dark for at least 7 days of analysis. As a control, *LacZ* was expressed using *tim*GAL4 and no effect on circadian activity profiles were observed. Circadian profiles in A-D are averaged here to 24 hour cycles for all the flies of each genotype, with variances plotted around the averaged profiles. (B) *tim*GAL4-driven knockdown of *Atxn7* causes major disruptions in circadian activity and causes a period lengthening. (C) Overexpression of wild-type *Atxn7* (*SCA7-10Q*) causes minimal effects on activity profiles. (D) Overexpressing polyglutamine-expanded *Atxn7* (*SCA7-100Q*) causes major disruptions in circadian activity profiles that closely phenocopy *Atxn7* knockdown. (E) Table of statistical analysis. Periods were determined by the peak of chi-square periodogram analysis for all rhythmic flies, and robustness of rhythmicity by Fast Fourier Transform (FFT) of all flies. Rhythmic flies produced a single robust peak with p<0.01 by chi-square periodogram analysis, while arrhythmic flies produced no statistically significant peak or multiple peaks. A nonparametric Kruskal Wallis ANOVA (p<0.0001) determined statistical significance of rhythmicity differences, while one-way ANOVAs showed statistically significant differences for FFTs (p<0.0001) and periods (p<0.0001).

The activity profiles from the *Atxn7* knockdown closely phenocopied that of Atxn7 poly-Q overexpression, and both showed alterations in period length and rhythmicity (Figure 7E). As a control, we expressed *LacZ* and did not observe the activity profile changes displayed in *Atxn7* RNAi flies (Figure 7A). These results demonstrate that *Atxn7* has profound consequences on circadian rhythm output. Average plots of activity for multiple flies for multiple days in constant darkness, plotted on a 24hr time scale are shown in Figure 7.

### Loss of *Atxn7* or *non-stop* differentially impact circadian clock gene expression

The *per* and *Clock* (*Clk*) genes are central components of the *Drosophila* molecular clock and they regulate rhythmic behaviors through transcriptional-translational feedback loops ((Abdalla, Mascarenhas, & Cheng, 2022; Fuhr et al., 2015; Hall, 2003; Koh, Zheng, & Sehgal, 2006; Luo et al., 2012; P. Meyer, Saez, & Young, 2006). Because these oscillations depend on precisely timed changes in mRNA levels, dynamics in *per* or *Clk* transcript rhythms provide a direct readout of how perturbing SAGA DUBm components alters the clock at a molecular level.

To assess whether loss of *Atxn7* or *non-stop* disrupts the circadian cycling of *per* or *Clk* transcripts, we used *tim*GAL4 to drive RNAi knockdown of each gene in clock cells and quantified mRNA expression across four circadian timepoints (ZT1, ZT7, ZT13, ZT19). Two-way ANOVA revealed highly significant effects of genotype, timepoint, and genotype-timepoint interaction on *per* transcript levels, indicating that both mean expression and circadian dynamics differ among *LacZ*, *Atxn7* RNAi, and *not* RNAi groups.

Individual Tukey post-hoc analysis showed that *LacZ*, *Atxn7,* and *non-stop* RNAi displayed differential cycling in *per* across ZTs (Figure 8A). Compared to controls, *Atxn7* and *non-stop* knockdown both displayed significant reductions in overall *per* transcripts relative to LacZ. *Atxn7* and *non-stop* RNAi also differed significantly from each other at ZT19, revealing a genotype-specific divergence at this phase of the cycle (Figure 8A). Together, these comparisons demonstrate that loss of either *Atxn7* or *non-stop* dampens *per* abundance and cycling, and *Atxn7* and *not* knockdown differ most strongly from each other late in the cycle. These results indicate that both genes are required to maintain normal *per* expression levels to preserve its circadian rhythm.

**Figure 8.**
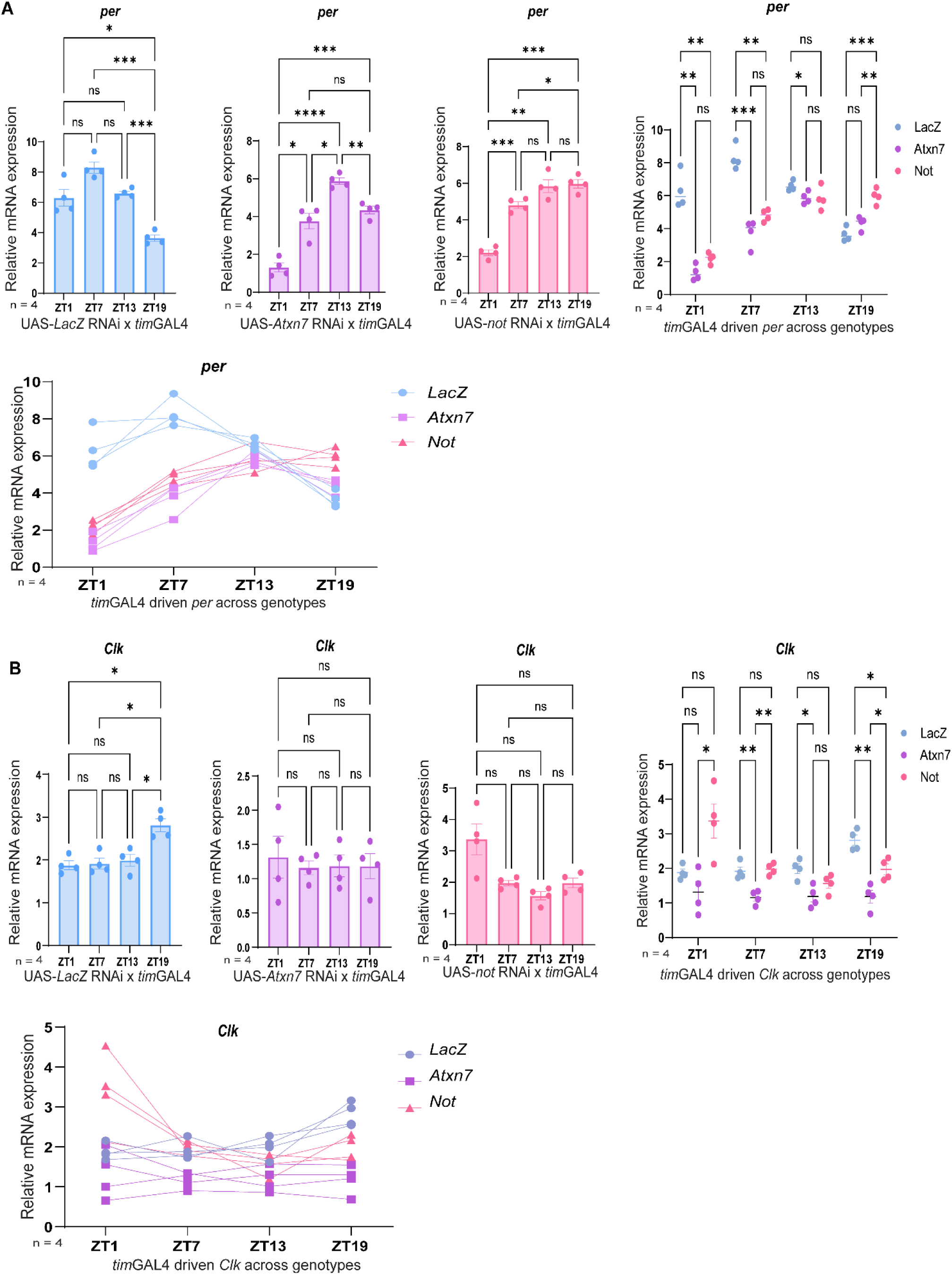
TimGAL4 driven knockdown of *Atxn7* and *not* in clock cells alters *per* and *Clk* gene expression. (A) Two-way ANOVA revealed significant effects of genotype (*p* < 0.0001) timepoint (*p* < 0.0001) and genotype-timepoint interaction (*p* < 0.0001) on *per* levels in the *tim*GAL4 driven RNAi’s. Tukey post-hoc tests showed significant but different circadian cycling in *per* across *LacZ*, *Atxn7* RNAi, and *non-stop* RNAi across ZTs, as seen in each of their individual graphs. The graph on the far right shows significant differences at each ZT amongst each genotype, and the graph below shows raw data points from all genotypes plotted together across circadian time. *p < 0.05, **p < 0.005, ***p < 0.0005, ****p < 0.0001, ns is not significant. Both *Atxn7* and *not* knockdown reduced *per* levels and altered cycling. (B) *Clock* transcript levels were affected by genotype (p < 0.0001), timepoint (*p* < 0.005) and their genotype-timepoint interaction (*p* < 0.0001). Post-hoc Tukey tests revealed no significant difference in *Clk* cycling between individual ZTs in *Atxn7* or *non-stop* RNAi but did show significance in *LacZ,* as seen in their individual graphs. *Atxn7* knockdown significantly reduced overall *Clock* mRNA relative to *LacZ* and *not* RNAi, whereas total *Clk* in *non-stop* knockdown did not differ from LacZ. The graph below shows raw data points from all genotypes plotted together across circadian time. *p < 0.05, **p < 0.005, ***p < 0.0005, ****p < 0.0001, ns is not significant.

Two-way ANOVA similarly demonstrated significant effects of genotype, timepoint, and their interaction on *Clk* mRNA expression. Post-hoc comparisons showed that *Atxn7* knockdown significantly reduced *Clk* transcript levels relative to both *LacZ* and *non-stop* RNAi (Figure 8B). In contrast, *non-stop* RNAi and *LacZ* did not significantly differ in total *Clock* mRNA levels between each other, but loss of *not* did disrupt *Clk* cycling. Thus, *Atxn7* and *non-stop* differentially effect *Clock* transcript levels and temporal regulation, with *Atxn7* showing more significance overall.

Taken together, these findings show that *Atxn7* and *non-stop* differentially regulate core clock gene expression. Both *Atxn7* and *non-stop* are required for proper *per* abundance and rhythmicity, with *non-stop* loss causing the most substantial disruptions in *per*. In contrast, *Clk* expression is particularly sensitive to *Atxn7* depletion, while *non-stop* primarily alters temporal patterning without affecting mean *Clk* transcript levels. This divergence highlights that distinct components of the SAGA DUBm influence circadian transcription through separable mechanisms and underscores the importance of Atxn7- and non-stop-dependent regulation in maintaining robust molecular clock function.

### *Atxn7* knockdown produces expression shifts without affecting total per protein levels

We next sought to determine whether regulation of SAGA DUBm on the circadian clock also occurs at a post-translational level. Knockdown of *Atxn7* in clock cells using *tim*GAL4 produced no changes on total per protein, although the peak of per expression was shifted from ZT19 consistently in wild-type, to ZT1 in the *Atxn7* knockdown, indicating a potential phase delay in per cycling (Figure 8B). The remaining ZTs in the *tim*GAL4-driven *Atxn7* RNAi retained a circadian pattern of expression similar to wild-type (Figure 8A,B).

### Atxn7 polyglutamine expansion in clock cells alters per protein expression

To assess whether polyglutamine-expanded *Atxn7* affects the circadian clock at a post-transcriptional level, flies overexpressing human *Ataxin-7-10Q* (*hAtxn7-10Q*) and -*92Q* (*hAtxn7-92Q*) in clock cells using *tim*GAL4 were entrained and heads collected across four ZTs.

Overexpression of *hAtxn7-10Q* did not significantly alter total per protein levels, however it demonstrated that peak per accumulation was shifted from ZT19 in wild-type to ZT1 in *hAtxn7-10Q* overexpression, with per signal approximately two-fold higher than wild-type at ZT1. The remaining ZTs in *hAtxn7-10Q* showed a similar circadian pattern and abundance to wild-type (Figure 8A,C,F,G).

Overexpression of h*Atxn7-92Q* caused a significant increase in total per protein, including elevated amounts of hyperphosphorylated per. Peak per expression also occurred at ZT1, with signal at ZT1 approximately five-fold higher than wild-type at that time point (Figure 8G). Per remained elevated throughout all timepoints in h*Atxn7-92Q* overexpression (Figure 8A,D).

Together with the qRT-PCR results, these findings demonstrate that Atxn7 is necessary for maintaining the proper regulation of the circadian clock at multiple levels.

**Figure 9.**
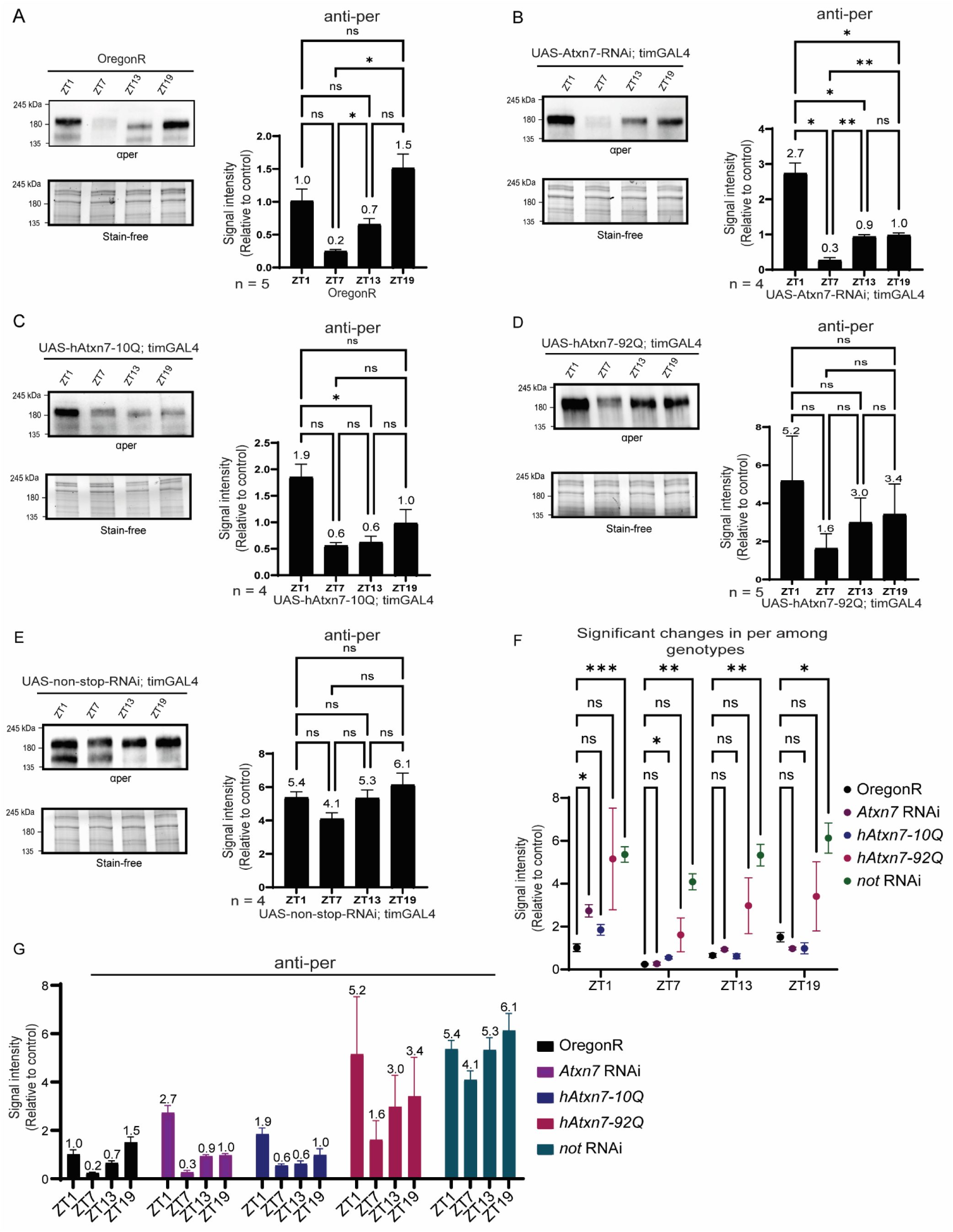
Mutations in Atxn7 and non-stop in clock cells disrupts per protein dynamics. (A) Western blot of endogenous per protein across four circadian timepoints (ZT1, ZT7, ZT13, ZT19) in wild-type flies, showing the expected circadian cycling in per. (B) *tim*GAL4-driven Atxn7 RNAi produces no major changes in total per abundance, but shifts the peak from ZT19 to ZT1. (C) overexpression of hAtxn7-10Q similarly preserves overall per levels but shifts peak accumulation to ZT1. (D) Overexpression of hAtxn7-92Q causes an increase in per levels and delays the peak accumulation to ZT1. (E) Non-stop RNAi produces the strongest per protein phenotype, with total per strongly elevated nearly five-fold or more at every ZT. (F) Quantification of per levels across genotypes showing significant differences among genotypes at each ZT (*p < 0.05, **p < 0.005, ***p < 0.0005, ****p < 0.0001, ns is not significant). (G) Quantification of per levels across genotypes plotted together. Error bars represent standard error. Two-way ANOVA was used for overall comparisons among all genotypes, and Brown-Forsythe with Welch ANOVA was used for ZT comparison within each genotype.

### *Non-stop* knockdown in clock cells causes substantial increases in period protein

Knockdown of *non-stop* gene in clock cells caused the most pronounced changes in *per* mRNA and protein dynamics. Total per protein levels displayed a striking five-and-a-half-fold elevation compared to wild-type, including the highest amounts of hyperphosphorylated or high molecular weight per out of any genotypes. Peak per occurred at ZT19, similar to wild-type, but overall per abundance and phosphorylation remained exceptionally elevated (4-6 times higher) at all timepoints across the circadian cycle (Figure A,E,G). Interestingly, per protein levels statistically did not significantly cycle throughout the day. As a deubiquitinase, our data suggest it is possible that non-stop regulates per stability at both the transcriptional and post-transcriptional level by controlling ubiquitin dynamics during its transcription and post-translationally controlling per degradation. Loss of *non-stop* may impair gene expression and per protein turnover, resulting in accumulation of per in its hyperphosphorylated, non-degraded form. Together, these results suggest the SAGA DUBm plays an important and necessary functional role in maintaining integrity of the circadian clock, through mechanisms at both the transcriptional and post-translational level.

## Discussion

Together, our findings demonstrate that the SAGA deubiquitinase module is interconnected to circadian regulation at multiple levels and is far more dynamic than previously appreciated. At the molecular level, both non-stop and Atxn7 exhibit dynamic oscillations in protein abundance across the circadian cycle. These rhythms are abolished in *per^0^* mutants, indicating that their temporal regulation depends on a functional circadian clock. Nonetheless, transcript levels for DUBm components in fly heads do not cycle, suggesting that the clock may exert control on the DUBm primarily through post-transcriptional or post-translational mechanisms.

In addition to abundance changes, we observe striking circadian dependent changes in the subcellular localization of non-stop and Atxn7, including brain regions where the two proteins do not co-localize. This spatial separation supports prior work showing that non-stop can dissociate from SAGA to form an independent DUB module (Lim et al., 2013). Our imaging results support a model in which the DUBm dynamically exists in SAGA-associated and independent states, with transitions in state and subcellular localization in a time-dependent manner. Such temporally controlled modularity aligns with earlier observations that non-stop participates in distinct biochemical pathways, such as stabilizing SCAR independently of SAGA, while regulating H2B ubiquitination as part of SAGA.

We also show that SCAR cycles in a circadian pattern, an effect lost in *per^0^* mutants. This rhythmicity further supports the idea that non-stops enzymatic activity is temporally tuned to coordinate both cytoskeletal and chromatin-associated processes. These results reinforce a broader model in which daily control of DUBm activity helps align cytoskeletal remodeling, transcriptional accessibility, and other ubiquitin-dependent pathways with circadian physiology.

To assess whether these DUBm components bidirectionally influence circadian output, we manipulated *Atxn7* and *non-stop* levels in the clock cells of the brain using *timeless-* and *Pdf*GAL drivers. Overexpression of wild-type *Atxn7* produced minimal effects on locomotor rhythms, whereas overexpression of polyglutamine-expanded *Atxn7* and RNAi-mediated knockdown of *Atxn7* each caused pronounced disruptions in circadian activity profiles, period length, and rhythmicity. These results indicate that both loss of *Atxn7* and disease-associated poly-Q expansion impair circadian activity rhythms, suggesting that proper Atxn7 dosage and integrity are required for stable rhythmic behavior.

We sought to determine at what molecular level the DUBm causes perturbations in the circadian clock and found it to be at both the transcriptional and post-transcriptional level. Knocking down *Atxn7* and *non-stop* caused significant alterations in *per* and *Clk* transcript cycling and abundance, along with alterations in per protein circadian cycling and abundance. Importantly, non-stop and Atxn7 displayed distinguishable effects on the dynamics of these clock components and also showed different effects in specific sets of clock cells.

*Non-stop* knockdown in clock cells using timGAL4 had the most substantial effects on both per transcript and protein dynamics. This suggests non-stop may have a role in maintaining proper per dynamics at both the transcriptional and post-transcriptional level. These findings point to a fundamental role for the SAGA DUBm in maintaining the molecular integrity of the circadian clock.

Taken together, our results support a model in which the SAGA deubiquitinase module functions as a dynamically regulated component of the circadian system, with rhythmic expression, localization, and substrate engagement. Perturbing this regulation, through loss of *not*, loss of *Atxn7*, or pathogenic polyglutamine expansion, disrupts clock dynamics and circadian behavior. These findings highlight the DUBm as a bidirectional player of the clock. Future work will be needed to define the upstream cues that drive rhythmicity and to determine how temporally dysregulated DUBm activity contributes to neurodegenerative phenotypes associated with Atxn7 dysfunction.

## Materials and Methods

### Immunoblot analysis

Protein samples were run on either 4-20% Biorad Stain-Free gradient gels (#4568093), or 8% polyacrylamide gels in 1X Laemmli electrode buffer. Stain-free polyacrylamide gels were imaged on a Chemidoc MP imaging system before transfer. Proteins were transferred to Amersham Hybond P 0.2 mM or 0.45 mM PVDF membranes (catalog number: 10600021 & 10600029) using the Trans-blot turbo transfer system from BioRad. The semi-dry transfer took place in Bjerrum Schafer-Nielsen Buffer with SDS (48 mM Tris, 39 mM Glycine, 20% Methanol, 0.1% SDS) at 1 Amp 25 V for 20 min. Membranes were then blocked for 1 hr in 5% non-fat dried milk diluted in 0.05% TWEEN-20 in 1XPBS. The membrane was then briefly rinsed and incubated in primary antibody diluted in 5% non-fat dried milk diluted in 0.05% TWEEN-20 for either 1 hr at room temperature or overnight at 4°C. The membrane was then washed four times for 5 min in 0.1% TWEEN-20 diluted in 1XPBS. The membrane was then incubated for 1 hr at room temperature in secondary antibody diluted 1:5000 or 1:10000 in 5% non-fat dried milk diluted in 0.05% Triton-X 100. The membrane was then washed four times for 5 min in 0.1% TWEEN-20 diluted in 1xPBS. The chemiluminescence was imaged on a Chemidoc MP Imaging system. Band intensities were measured on ImageLab software.

### Immunofluorescence

Drosophila adult or 3^rd^ instar larval brains were dissected into 4% methanol free formaldehyde/1xPBS and fixed at 4°C for 1 hr. Fixative was removed and the brains were washed with 1xPBS and then either stored at 4°C in 1XPBS for up to 1 week or immediately stained. Brains were washed three times in 1xPBS containing 5% Triton X-100. They were then blocked in 0.2% TWEEN 20% and 5% BSA diluted in 1xPBS for 1 hour at room temperature.

Primary antibodies were incubated for 1-2 nights and then washed four times 10 min at room temperature in 1xPBS containing 5% Triton X-100. Brains were incubated in Alexafluor conjugated secondary antibodies diluted 1:1000 in blocking buffer overnight at 4°C. They were then washed four times for 10 min at room temperature in 1XPBS containing 5% Triton X-100. Brains were mounted in Vectashield containing DAPI (Vector Labs catalog number: H-1200).

### Antibodies

Mouse anti-SCAR (DSHB P1C1-SCAR RRID:AB_2618386) was used at a dilution of 1:300 for immunoblotting and 1:50 for immunofluorescence. Guinea pig anti-non-stop (Mohan et al., 2014) was used at a dilution of 1:2000 for immunoblotting and 1:XXX for immunofluorescence. Rabbit anti-Atxn7 (Mohan et al., 2014) was used at a dilution of 1:5000 for immunoblotting and 1:1000 for immunofluorescence. Mouse anti-Ubiquitinated H2B antibody was purchased from Millipore (05-1312) and used at a dilution of 1:4000. Goat anti-Guinea Pig HRP (Jackson ImmunoResearch INC catalog number: 106-035-003, RRID:AB_2337402) was used at a dilution of 1:10,000. Goat anti-mouse HRP (Polyclonal) (Jackson ImmunoResearch INC catalog number: 115-035-003, RRID:AB_10015289) was used at a dilution of 1:5000. Goat anti-Rabbit HRP (Polyclonal) (Jackson ImmunoResearch INC catalog number: 111-035-003, RRID:AB_2313567) was used at a dilution of 1:10,000. The rabbit anti-period antibody was obtained as a gift from Jeffery Price and was used at a dilution of 1:20000. For immunofluorescence, Goat anti-Rabbit and IGG-488 (Invitrogen Catalog number: A-11008) was used at a dilution of 1:1000. Goat anti-Guinea pig IGG-568 (Invitrogen Catalog number: A11075) was used at a dilution of 1:1000. Goat anti-Mouse IGG-594 (Invitrogen Catalog number: A11005) was used at a dilution of 1:1000.

### Protein Sample Preparation

Flies were collected in liquid nitrogen at ZT1, 7, 13, and 19 and stored at -80 Celsius until analyzed. Heads were separated using liquid nitrogen or dry ice and homogenized in 2X UREA sample buffer. For each head, 7ul of sample buffer was added. Extracts were boiled at 80°C for 10 min and spun down at 13000rpm for 7 min. Samples were ran immediately or stored in 20°C. For experiments using non-stop mutants, larval brains were used and dissected into 1xPBS. The PBS was removed and 2xUREA sample buffer (50 mM Tris HCl pH 6.8, 1.6% SDS, 7% glycerol, 8M Urea, 4% b-mercaptoethanol, 0.016% bromophenol blue) was added in a 1:1 ratio and 10 units of Benzonase were added. The brains were homogenized by pipetting and boiled at 80°C for 10 min and spun down at 13000rpm for 7 min.

### qRT-PCR

Total RNA was extracted by using TRIzol reagent (Invitrogen, Waltham, MA, USA). Extracted RNA was treated with TURBO DNAse (Invitrogen, Waltham, MA, USA) to remove contaminating DNA. Reverse transcription was performed with the High-Capacity cDNA Reverse Transcription Kit (ABI, Waltham, MA, USA). Messenger RNA levels were quantified by using the StepOnePlus Real-Time PCR System with Fast SYBR Green Master Mix (ABI, Waltham, MA, USA). Rp49 was used as internal control. At least three biological replicates were used. Statistical analysis was performed in GraphPad Prism and specified in figure legends. Primers used to detect the transcripts are shown in table 1.

**Table 1.**
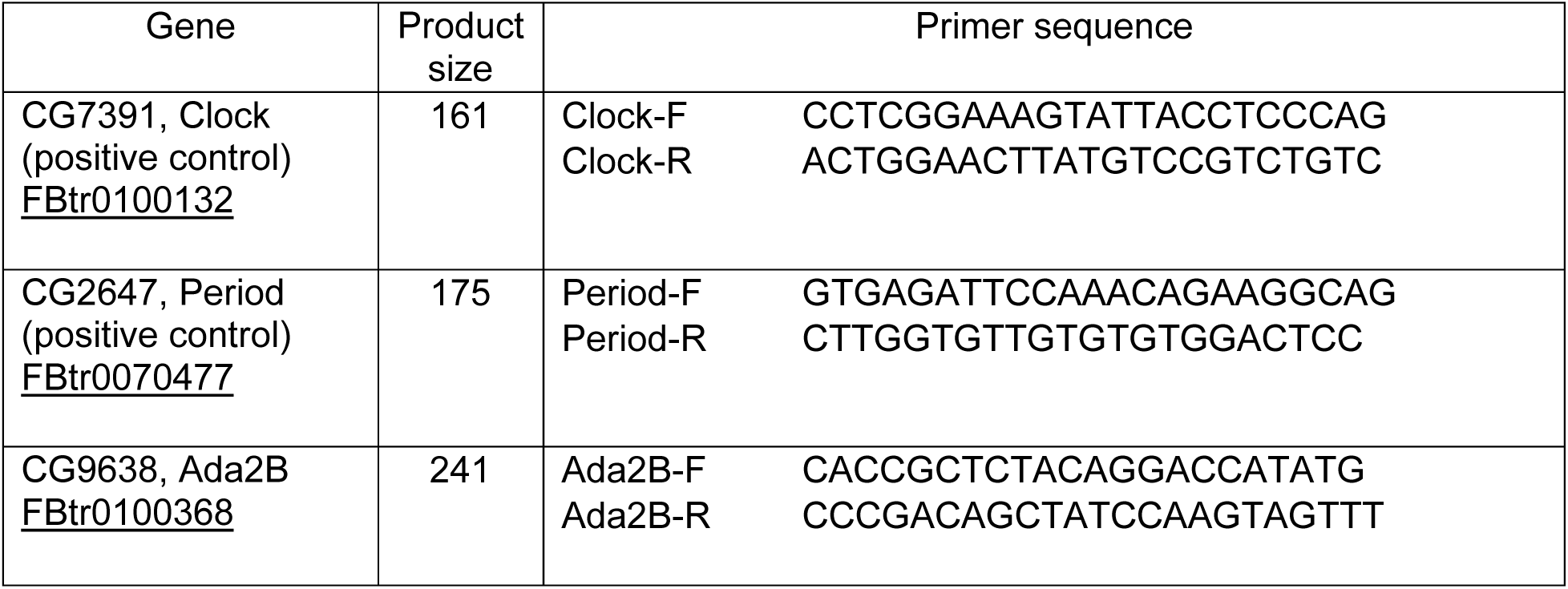
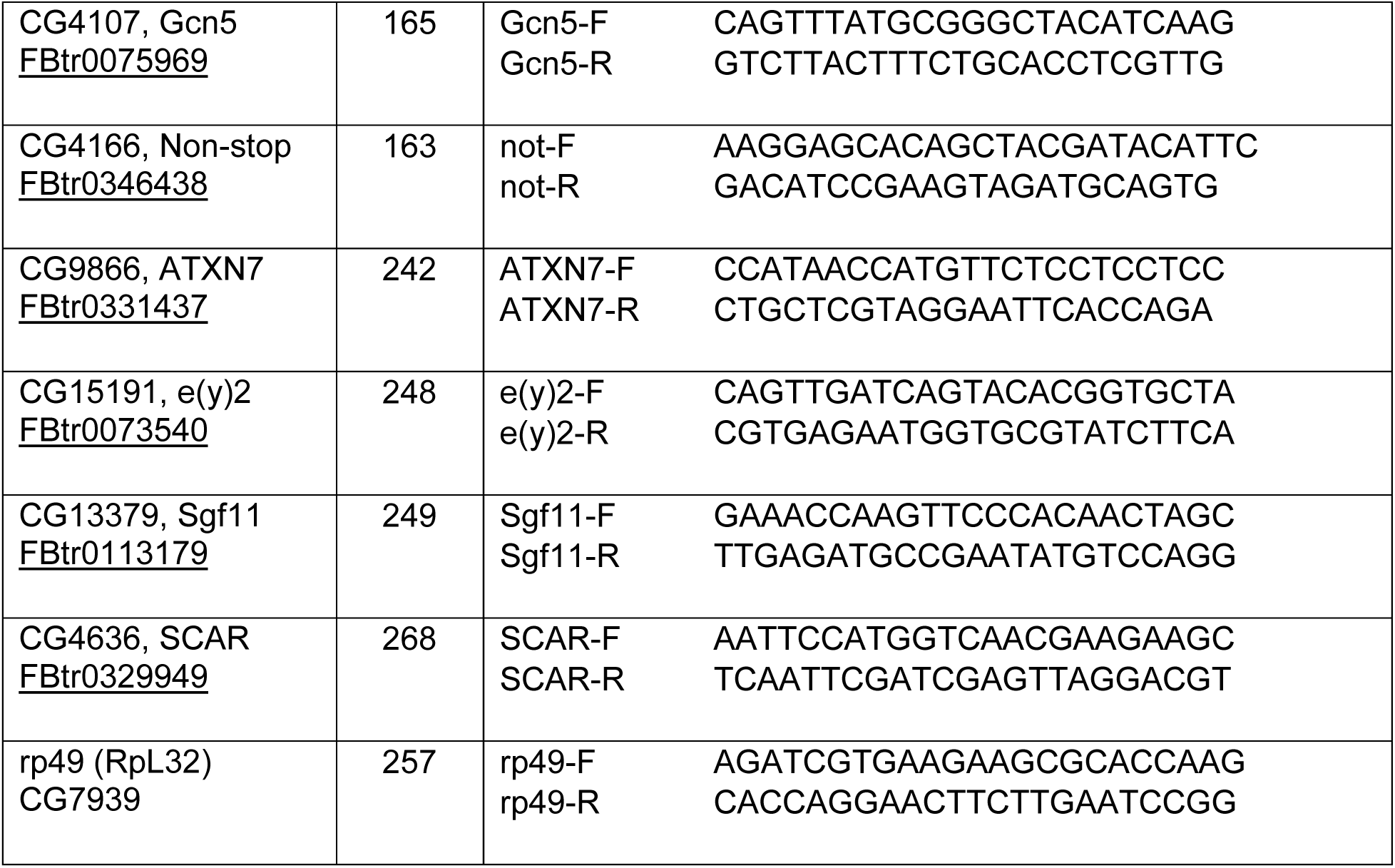
Primers generated for qRT-PCR.

### Fly Entrainment and Locomotor Assays

The RNAi lines were crossed to a *tim*GAL4 and *Pdf*GAL4 driver, limiting expression to clock-specific neurons because the GAL4 expression was dependent on the activity of the *timeless* and *Pdf* promoters. The crosses were entrained at 23.5 °C under a 12-h/12-h light-dark (LD) cycle with cool white fluorescent bulbs (ca 3000 lx). The F1 progeny containing the *tim*GAL4/*Pdf*GAL4 driver and the UAS-*RNAi* responder were collected and entrained for a further 72 h in separate vials at 23.5 °C. Males were used for locomotor assays (because of their more robust locomotor activity). The male flies were loaded into individual cuvettes and placed in a monitoring device connected to a computer (Trikinetics, Waltham, MA), and data were recorded and analyzed using the method described previously by Muskus et al. 2007.

### Analysis of Circadian Locomotor Activity Rhythms

UAS-*Atxn7* RNAi and UAS-*not* RNAi lines were crossed with UAS-*dcr2;tim*GAL4 and UAS-*dcr2*;*Pdf*GAL4 lines. Male flies placed into glass cuvettes were assayed in behavioral monitors from Trikinetics (Waltham, MA, USA), with 10 min collection bins. The flies were entrained for at least three days to LD 12 h:12 h and then monitored in constant darkness for 7 or more days.

Individual actograms were obtained for all flies, and these were then averaged with a 24 hour period for all flies of a given genotype. ClockLab was used to perform chi-square periodogram analysis to determine rhythmicity and period length. Rhythmic flies produced single statistically significant peaks, and the period length of the peak was taken as the period for each fly. Rhythm intensity was the average Fast Fourier Transform for each genotype.

### Fly stocks

The following fly lines were used in this study. Wild-type Oregon-R (DGGR Catalog number: 109162, RRID:DGGR_109612), the *non-stop^02069^* line P(ry[+t7.2]=PZ)not[02069] ry[506]/TM6B, ry[CB] Tb[+] was obtained from the Bloomington *Drosophila* Stock Center (BL115533). The *per^0^* fly line, y[1] per[01] w[*] was obtained from the Bloomington *Drosophila* Stock Center (BL80917). *Non-stop* RNAi (#1) (y[1] v[1]; P{y[+t7.7] v[+t1.8]=TRiP.JF03152}attP2/TM3, Sb[1]) was obtained from Bloomington *Drosophila* Stock Center (BL28725). *Non-stop* RNAi (#2) (y[1] v[1]; P{y[+t7.7] v[+t1.8]=TRiP.JF03152}attP2/TM6B, Tb) was rebalanced from Bloomington *Drosophila* Stock BL28725. UAS-Sgf11 RNAi was obtained from the Vienna *Drosophila* Resource Center VDRC:100581. UAS-*dcr2;tim*GAL4 and UAS-*dcr2*;*Pdf*GAL4 were gifts from Jeffrey Price. UAS-*Atxn7* RNAi (P KK110634 VIE-260B) was obtained from the Vienna *Drosophila* Resource Center VDRC:102078 RRID:FlyBase_FBst0473949, Construct ID:110634. UAS-*not* RNAi was obtained from the Vienna *Drosophila* Resource Center stock #45776.

### Statistics

All other statistical tests were conducted using Prism 9 (GraphPad) and are specified in figure legends. The number of biological replicates is noted on figures and corresponding legends.

## Acknowledgements

We thank the Bloomington Drosophila Stock Center for stocks obtained in this study. We thank Dr. Jeffrey Price for his collaboration and providing the period antibody and several fly lines. We thank all current and past Mohan laboratory members for helpful comments and suggestions.

